# Light-sheet imaging and graph analysis of antidepressant action in the larval zebrafish brain network

**DOI:** 10.1101/618843

**Authors:** Jessica Burgstaller, Elena Hindinger, Joseph Donovan, Marco Dal Maschio, Andreas M. Kist, Benno Gesierich, Ruben Portugues, Herwig Baier

**Author notes:** Correspondence concerning this article should be addressed to H. B.

## Abstract

The zebrafish is increasingly being employed as an experimental platform to model neuropsychiatric diseases and to screen for novel neuro-active compounds. While the superb genetic and optical access that this system offers has long been recognized, these features have not been fully exploited to investigate disease mechanisms and possible therapeutic interventions. Here we introduce a light-sheet imaging and graph-theoretical analysis pipeline to determine the effects of the known or suspected antidepressant compounds fluoxetine, ketamine and cycloserine on brain-wide neural activity patterns. We imaged the brains of both wildtype fish and *gr^s357^* mutants, which harbor a missense mutation that abolishes glucocorticoid receptor transcriptional activity. The *gr^s357^* mutation results in a chronically elevated stress axis together with behavioral endophenotypes of depression. Consistent with broad expression of the glucocorticoid receptor throughout the brain, we show that the mutant fish exhibit an altered correlational structure of resting-state brain activity. Intriguingly, in *gr^s357^* mutant fish, an increased ‘modularity’, which represents the degree of segregation of the network into highly clustered modules, was restored by acute fluoxetine administration to wildtype levels. Ketamine and cycloserine also normalized specific parameters of the graph. Fluoxetine altered network function in the same direction in mutant and wildtype, while ketamine and cycloserine had effects that were opposite for the two genotypes. We propose that light-sheet imaging, followed by graph analysis, is a content-rich and scalable first-pass approach for studying the neural consequences of drug effects and drug x genotype interactions in zebrafish models of psychiatric disorders.

## Introduction

Recent work has highlighted the promise of the zebrafish model to understand mutations in psychiatric disease genes and drug actions (for a review, see Haesemeyer & Schier, 2015). Zebrafish larvae carrying a mutation of the GR (*gr^s357^*) have a chronically elevated hypothalamic-pituitary-adrenal (HPA) axis due to disruption of negative feedback via cortisol (Fig. 1a). The *gr^s357^* mutants swim less and startle more than wildtype siblings at larval stages. When repeatedly placed into a novel tank, adult mutants fail to habituate and rather exaggerate their behavioral response (‘freezing’) to this mild stressor. Importantly, their HPA activation and behavior can be normalized by acute or chronic treatment with the selective serotonin reuptake inhibitor (SSRI) fluoxetine (Griffith et al., 2012; Ziv et al., 2013). Moreover, it has been shown that reducing HPA axis activity and enhancing serotonergic transmission via treatment with fluoxetine shifts neuronal responses and processing of food stimuli in the fish (Filosa et al., 2016). One of our goals therefore was to evaluate the potential of fluoxetine to restore altered brain function.

**Figure 1.**
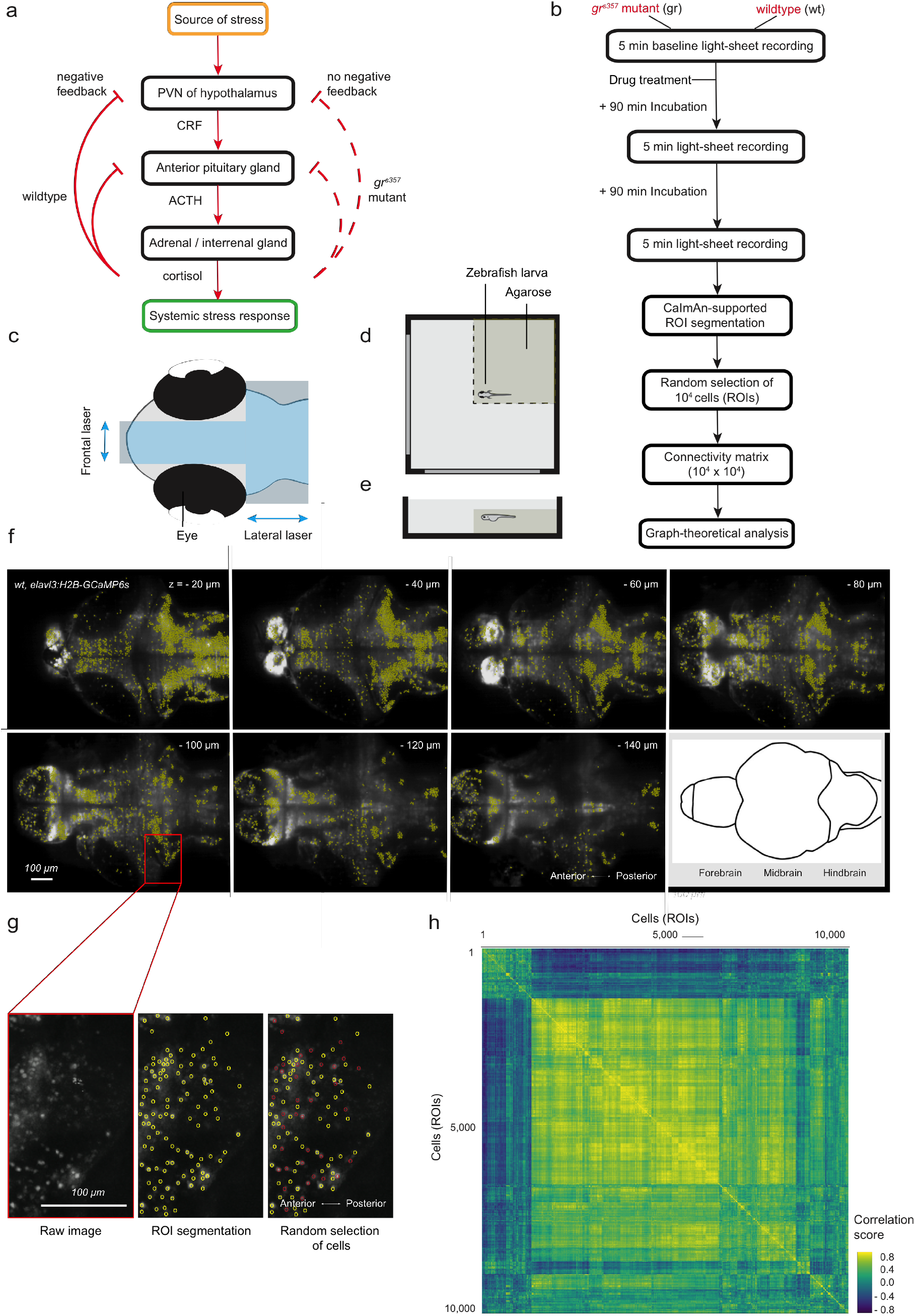
Experimental workflow. **(a)** Dysregulated HPA axis in *gr^s357^* zebrafish mutants. The *gr^s357^* mutation abolishes glucocorticoid receptor transcriptional activity and thus negative feedback on the stress response, which results in a chronically elevated stress axis together with endophenotypes of depression. **(b)** Experimental workflow. The first light-sheet recording before drug application informs about baseline brain activity of the two genotypes tested (*gr^s357^* mutant, wildtype fish). Thereafter, fish are incubated in the setup for 1.5h with fluoxetine (20μM), ketamine (20μM), or cycloserine (20μM). DMSO serves as vehicle control. Between the second and third recording, fish stay in the setup and incubate for another 1.5h. Graph-theoretical analysis is applied to all three recordings: 0, 1.5h and 3h drug treatment. **(c)** Sample preparation in the light-sheet microscope. Laser beam cross (in blue) on head of larva. **(d)** Top view of embedded larva in a custom chamber. Yellow indicates a block of agarose cut along dashed lines. **(e)** Side view of larva in chamber. **(f)** CaImAn-supported ROI detection of neurons in seven representative planes of 30 planes acquired per brain. **(g)** Close up of cellular resolution before (left) and after ROI detection (middle), and the randomly selected subsample of ROIs in red for further analysis (right). **(h)** Hierarchical clustering of functional connectivity. Example of a neuron-to-neuron correlation matrix for fluorescence traces of 10^4^ randomly selected neurons in one representative fish, as used for network construction and graph analysis of functional connectivity. Neurons were sorted by hierarchical clustering.

In addition to fluoxetine, we investigated ketamine and cycloserine, two potential antidepressant compounds with different mechanisms of action. Ketamine is a rapid-acting N-methyl-D-aspartate (NMDA) receptor antagonist and has recently been approved by the US Food and Drug Administration (FDA) as a nasal spray for the treatment of SSRI-resistant depression. Ketamine has also been shown to modulate zebrafish behavior (Michelotti et al., 2018), although its range of actions in this model system is poorly understood. Cycloserine is a natural antibiotic product of *Streptomyces orchidaceus* and *Streptomyces garyphalus* and has been employed in tuberculosis therapy since the late 1950s (Offe, 1988). Years later it was postulated, and later proven in slice preparations, that the compound influences long-term potentiation, a neuronal mechanism thought to be relevant for learning (Watanabe et al., 1992). Cycloserine is a partial NMDA agonist (Watson et al., 1990); in vivo, it acts like an agonist at low doses, but has antagonistic features at high doses. Its potential for the treatment of neurological and psychiatric conditions, such as major depression, is currently being investigated.

Here we used light-sheet calcium imaging for recording of spontaneous brain-wide activity in zebrafish (Ahrens et al., 2013) and analyzed the correlational structure of the data with a computational platform based on graph theory. This approach was inspired by work in human functional magnetic resonance imaging (Ye et al., 2015), which employs voxel-based graphs to reveal global patterns of brain activity. Recording of tens of thousands of single-cell activity traces generates a deluge of data, and we reasoned that graph analysis offers a relatively straightforward method to extract informative statistical parameters from the resulting adjacency matrix. Using a similar graph-based approach, albeit with far fewer nodes, Avitan et al. (2017) studied the developmental reorganization of spontaneous activity in the tectum. We compared whole-brain functional connectivity in wildtype and *gr^s357^* mutants and detected genotype-linked differences. Finally, we showed that, in line with normalizing *gr^s357^* behavior (Griffith et al., 2012; Ziv et al., 2013), acute fluoxetine administration restored disrupted network parameters in the mutant fish. Ketamine and cycloserine also differentially affected specific parameters of whole-brain functional connectivity. Our experimental workflow, light-sheet imaging followed by first-pass graph analysis, represents a content-rich, sensitive and scalable method to explore genetic and pharmacological alterations of brain-wide network activity in larval zebrafish.

## Results and Discussion

### Light-sheet imaging of resting-state brain activity in larval zebrafish

To record spontaneous (resting state) whole-brain activity, we embedded mutant and wildtype zebrafish larvae, at 6 days post fertilization (dpf), in low-melting agarose gel. The animals carried the transgene *Tg*(*elavl3:H2B-GCaMP6s*), encoding a nuclear-localized GCaMP6s under the control of a pan-neuronal promoter (Fig. 1d, e). For each condition, eight *gr^s357^* mutants and eight wildtype fish were imaged at cellular-resolution with a custom-built light-sheet microscope, as described in Methods. Data were acquired in 5 min whole-brain recordings at 2 Hz, resulting in 600 scanned volumes at baseline and after incubation with fluoxetine (20μM), ketamine (20μM) and cycloserine (20μM) for 1.5h and 3h in each fish (Fig. 1b). The concentration of 20μM was chosen to approximate the conditions of small-molecule compound screens in zebrafish larvae (Rihel et al., 2010; Kokel & Peterson, 2011). A recording time of 5 min was chosen to simulate measurements of resting-state brain activity by fMRI for functional connectivity analysis in human subjects with major depression (Greicius et al., 2007). We reasoned that this time window should be long enough to capture the normally occurring cycles of spontaneous brain activity in zebrafish. Although not tested systematically, drug incubation longer than 3h, or more frequent recordings within the 3 hour time window, appeared to result in inconsistent results, probably caused by stress, drug toxicity, photo-damage in the light-sheet microscope, or combinations thereof. Examples of motion-corrected, but otherwise raw fluorescence signals from single imaging planes are provided in two supplementary movies (one mutant, one wildtype).

### Segmentation of brain-wide functional imaging data at single-cell resolution

To examine functional connectivity, four-dimensional imaging stacks (time, x, y, z) were split into time series of single planes and corrected for motion artefacts using a customized version of the CaImAn package (Pnevmatikakis et al., 2017; Giovannucci et al., 2019) (see Methods). Regions of interest (ROIs) were spatial clusters of co-active pixels, likely corresponding to single neurons (Fig. 1f, g) (Portugues et al., 2014). The CaImAn algorithm picked up between 20,000 and 40,000 cells for each fish brain, i. e., ca 25% of all neurons. A subset of them, 10,000 ROIs were selected randomly for further analysis of network properties (Fig. 1g). This reduction in the number of cells analyzed reduced the computational load, and thus CPU time, generated by the high-dimensional connectivity matrix by two orders of magnitude (100 Million vs. 1600 Million matrix components) without noticeably affecting the statistics of the overall network structure. Functional connectivity was computed as Pearson’s correlation coefficients between ΔF/F signal time series of these ROIs (Fig. 1h).

### Analysis of functional imaging data by graph analysis

A powerful way to describe the functional organization of the whole brain is to treat the brain network as a graph. A graph is defined as a mathematical object consisting of a set of items, called nodes, and pairwise relationships between them, called edges (Bollobas, 1998). In our datasets, nodes were defined as the 10^4^ randomly selected cells (ROIs), and edges were defined by calculating Pearson correlation coefficients between ROI signal time courses. Hence, the entirety of all edges was represented by a 10^4^ × 10^4^ correlation matrix (Fig. 1h). Because edges from a node to itself are not allowed in simple graphs, the diagonal of each matrix was set to zero and the matrix was thresholded at 0.25, in order to ignore negative edges and remove spurious connections. Interactions between network nodes are best described by weighted connections, where the weight indicates the score of correlation and strength of interaction between nodes. Thus, a weighted connectivity matrix was derived to extract neural assemblies from the recordings (see Methods).

Many studies have focused on local properties of networks, such as clustering (Watts & Strogatz, 1998), degree distributions (Barabasi & Albert, 1999; Amaral et al., 2000) and correlations (Pastor-Satorras et al., 2001; Newman, 2002). There have also been studies that examine large-scale properties such as path lengths (Watts & Strogatz, 1998), percolation (Cohen et al., 2000; Callaway et al., 2000) or hierarchy (Ravasz & Barabasi, 2003; Clauset et el., 2008). For our study, we chose the following standard metrics (Box 1): 1) clustering coefficient, 2) characteristic path length, 3) modularity and 4) small-worldness.

#### Box 1. Standard metrics for analysis of whole-brain functional connectivity graphs.

##### Clustering coefficient

The clustering coefficient *C* describes the fraction of a node’s neighbors that are neighbors of each other. The global level of clustering *C_g_* in a network is the average of the local clustering coefficients of all the nodes *n*.

Clustering on a neuronal level is interpreted as a measure of local efficiency of information transfer.

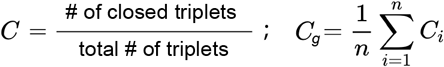

##### Modularity

Modularity describes a subdivision of the network into groups of nodes in a way that maximizes the number of within-group edges, and minimizes the number of between-group edges. This metric quantifies the degree to which the network may be subdivided into such groups.

Modules in a neuronal network might correspond to functional circuits that perform certain tasks.

##### Characteristic path length

The characteristic path length *l_G_* is the minimum number of edges between two nodes *v*_1_, *v*_2_ in the network. It is a measure of functional integration with shorter paths implying stronger potential for integration. In the brain network it is related to the capability for parallel information propagation.

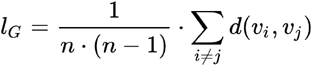

##### Small-worldness

Small-worldness is a property of networks that are highly clustered, like regular lattices, yet have small characteristic path lengths, like random graphs. Small-world networks possess high local, as well as global, efficiency in signal processing. In our dataset, we calculated small-worldness *ω* as the fraction of the mean clustering coefficient and the mean characteristic path length of the graph. The functional organization of the brain appears to be designed with such small-world network characteristics, because it allows both highly efficient local information processing in specialized modules and global integration at a relatively low wiring cost.

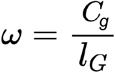

### Differences in functional connectivity between *gr^s357^* mutants and wildtype

We explored genotype differences in the dataset after incubation with DMSO in the control condition (Fig. 3). Due to deviations of the data from a normal distribution according to Shapiro Wilk test statistics, non-parametric testing with the Mann-Whitney U test was performed and the alpha-level was Bonferroni corrected according to multiple comparisons. The clustering coefficient (Fig. 3a), characteristic path length (Fig. 3c) and small-worldness (Fig. 3d) were not significantly different between *gr^s357^* mutant and wildtype fish at baseline (*p* > 0.05). However, modularity (Fig. 3b) was significantly higher in the *gr^s357^* mutant fish (*p* = 0.007). Mapping back the ROIs of detected modules to individual fish brains revealed that the number of modules in *gr^s357^* mutant fish was significantly greater compared to wildtype fish at baseline (Fig. 2a, b) (*p* = 0.034). These results suggest that the functional organization of the mutant brain comprises a greater number of relatively more isolated communities of neurons that seem to be diversely distributed across the brain (Fig. 2b). In wildtype fish the modules seem to be rather separated and located in distinct regions, with the forebrain comprising one module and a second module being located in the hindbrain (Fig. 2a). After 1.5h in the setup and incubation with DMSO, the clustering coefficient was significantly lower in the *gr^s357^* mutant compared to the wildtype fish (*p* = 0.005). The same holds true for small-worldness in the brain network (*p* = 0.001). The characteristic path length was significantly higher in the *gr^s357^* mutant fish compared to wildtype (*p* = 0.029). After incubation for 3h, genotype differences were present in all variables. Clustering and small-worldness were significantly lower in the *gr^s357^* mutant fish compared to wildtype (*p* = 0.001; *p* = 0.000), while characteristic path length and modularity were significantly (*p* = 0.011; *p* = 0.008) increased. These results suggest that time spent in the setup seems to differentially affect mutant and wildtype fish. Whilst the mutant fish have a chronically elevated stress level, wildtype fish might naturally respond to the stress experienced during the experimental procedure, which in turn affect network properties. A change in network parameters observed in wildtype, while absent in mutants, may imply a ceiling effect of the stress axis on functional connectivity in the *gr^s357^* brain.

**Figure 2.**
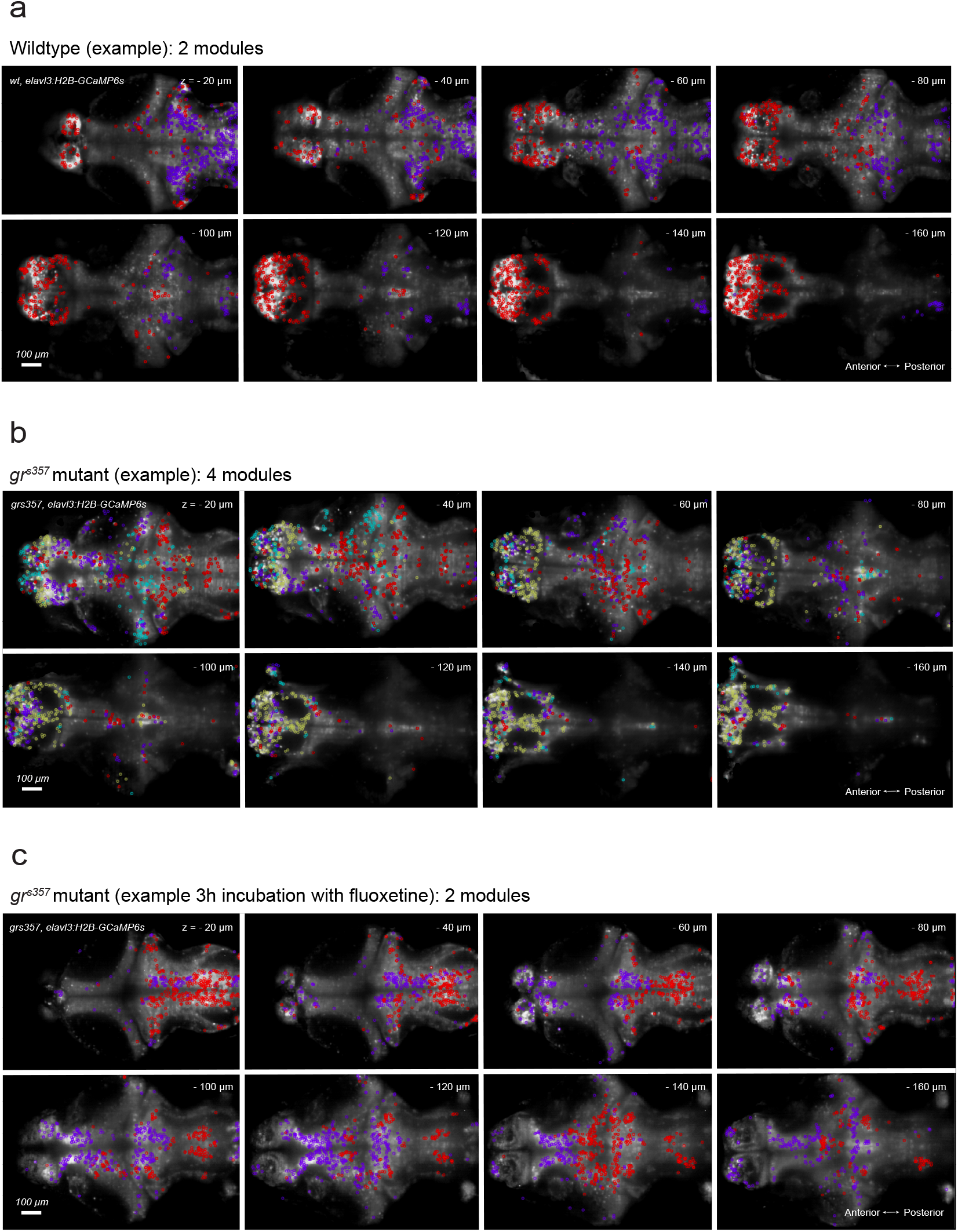
Representative examples of the modular organization of functional brain networks. **(a)** Example of a wildtype fish with 2 modules. **(b)** Example of a *gr^s357^* mutant fish with 4 modules. **(c)** Example of a *gr^s357^* mutant fish with 2 modules after 3h incubation with fluoxetine.

**Figure 3.**
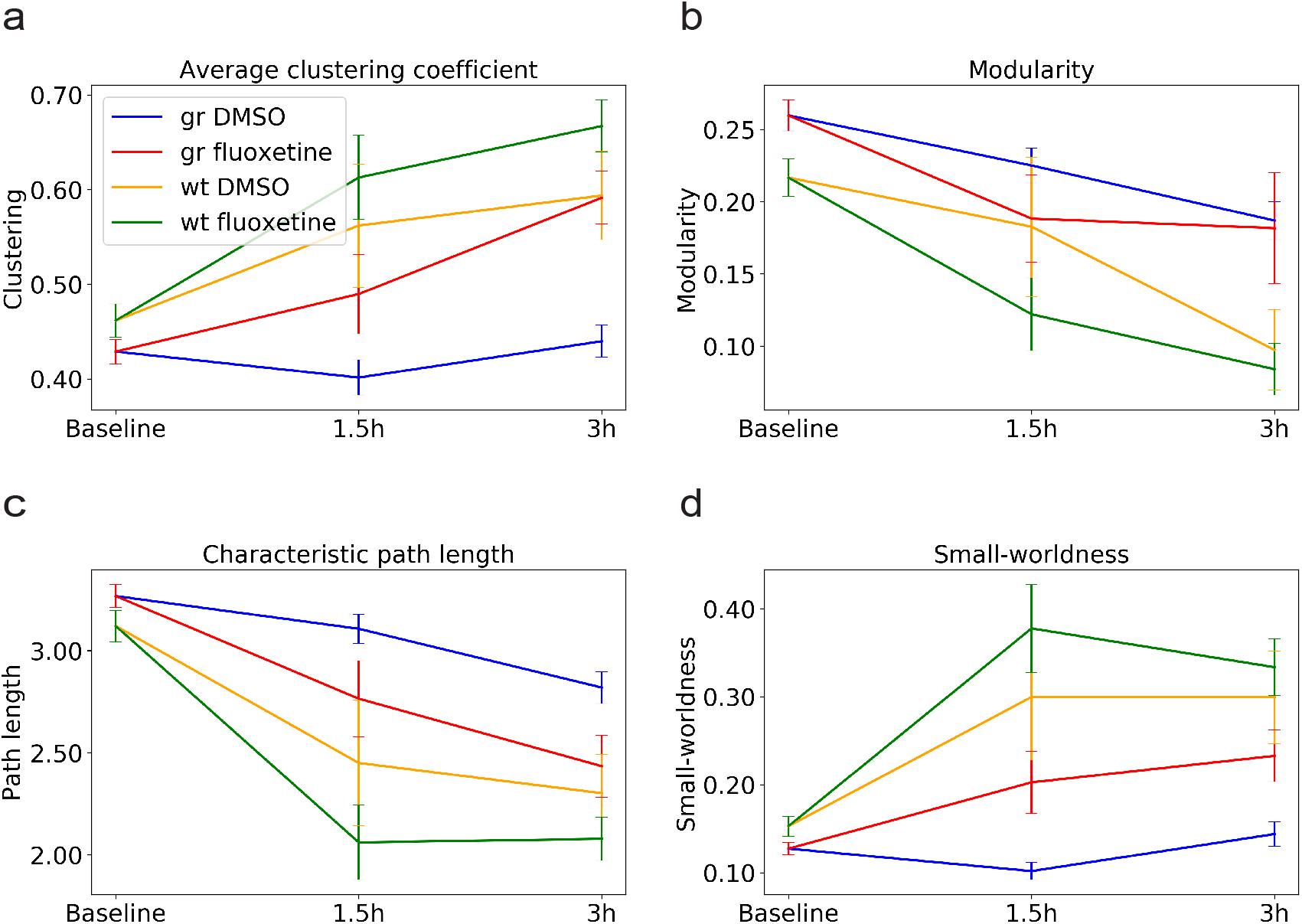
Effect of genotype and drug treatment with fluoxetine on network parameters. Mean of the **(a)** clustering coefficient, **(b)** modularity, **(c)** characteristic path length and **(d)** small-worldness in *gr^s357^* mutant and wildtype (wt) fish before drug treatment (baseline) and after 1.5h and 3h incubation with fluoxetine (20μM) or DMSO. 8 fish were imaged for each combination of genotype (*gr^s357^*, wt) and treatment (fluoxetine, control). Error bars represent the standard error of the mean.

### Normalization of brain-wide connectivity in the *gr^s357^* mutant by fluoxetine

We hypothesized that acute fluoxetine administration might restore network connectivity in *gr^s357^* fish. Indeed, we observed that clustering (*p* = 0.125), characteristic path length (*p* = 0.171), modularity (*p* = 0.476) and small-worldness (*p* = 0.076) were no longer significantly different between fluoxetine-treated mutant and (DMSO-treated) wildtype brains. The same holds true for incubation over 3h, where there is no difference in clustering (*p* = 0.492), characteristic path length (*p* = 0.303), modularity (*p* = 0.090), number of modules (*p* = 0.473) (Fig. 2c) and small-worldness (*p* = 0.235). We propose that acute fluoxetine treatment has the potential to restore whole-brain functional organization of the mutant fish brain, an effect that may explain the normalizing effect on behavior observed in our earlier study (Griffiths et al., 2012).

When comparing drug effects, we observed that fluoxetine affects all network parameters, albeit not all of them with statistical significance, in the same direction for both genotypes. Incubation with fluoxetine for 3h increased clustering in mutants and wildtype (Fig. 3a; GR DMSO vs. GR fluoxetine: *p* = 0.000; WT DMSO vs. WT fluoxetine: *p* = 0.085), decreased characteristic path length (Fig. 3c; GR DMSO vs. GR fluoxetine: *p* = 0.016; WT DMSO vs. WT fluoxetine: *p* = 0.091), and increased small-worldness (Fig. 3d; GR DMSO vs. GR fluoxetine: *p* = 0.001; WT DMSO vs. WT fluoxetine: *p* = 0.165). Modularity was not significantly different between genotypes after 3h incubation (Fig. 3b; GR DMSO vs. GR fluoxetine: *p* = 0.197; WT DMSO vs. WT fluoxetine: *p* = 0.165). We conclude that there is little, if any, drug x genotype interaction (Table 1), and the direction of fluoxetine’s effect on whole-brain functional connectivity is independent of the animal’s chronic stress level.

**Table 1.** Drug-induced changes in *gr^s357^* mutant (left arrow in blue) and wildtype (right arrow in red) network parameters. Compounds that differentially affected *gr^s357^* mutant and wildtype fish after 1.5h or 3h incubation are highlighted. 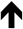 indicates an increase in the graph metric; 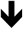 indicates a decrease; and – indicates no change compared to the genotype-specific DMSO control. Dark arrows indicate statistical significance; light arrows indicate a non-significant trend. Solid green filling indicates a double-sided significant drug x genotype interaction; hatched green indicates a significant interaction in one direction.

|  | Fluoxetine<br>1.5h | Fluoxetine<br>3h | Ketamine<br>1.5h | Ketamine<br>3h | Cycloserine<br>1.5h | Cycloserine<br>3h |
| --- | --- | --- | --- | --- | --- | --- |
| Clustering<br>coefficient | ↑↑ | ↑↑ | ↓↓ | ↑↓ | ↑↓ | ↓↓ |
| Modularity | ↓↓ | ↓↓ | ↑↑ | ↓↑ | ↓↓ | –↑ |
| Characteristic<br>path length | ↓↓ | ↓↓ | ↑↑ | ↓↑ | ↓↑ | –↑ |
| Small-worldness | ↑↑ | ↑↑ | ↑↓ | ↑↓ | ↑↓ | –↓ |

### Restoration of clustering in *gr^s357^* mutants by ketamine

After 1.5h of incubation with ketamine (20μM), clustering (Fig. 4a) was significantly reduced in *gr^s357^* mutants compared to wildtype control (*p* = 0.002). The characteristic path length (Fig. 4c) was increased (*p* = 0.005), modularity (Fig. 4b) was not significantly different (*p* = 0.109), and small-worldness (Fig. 4d) was reduced (*p* = 0.002). After incubation for 3h, there was no difference in clustering (*p* = 0.166), but modularity and characteristic path length were significantly higher in the ketamine treated *gr^s357^* fish compared to wildtype control (*p* = 0.038; *p* = 0.047). Smallworldness was significantly reduced in the mutant brain network (*p* = 0.033). These results suggest that clustering is the network parameter most affected by treatment with ketamine. Since brain networks of ketamine-treated *gr^s357^* mutants show similar clustering as wildtype, we conclude that ketamine reduces segregation and thus enhances the efficiency of local information transfer in the mutant brain. This may be the functional correlate of reduced inhibition in the brain by ketamine acting as an NMDA receptor antagonist.

**Figure 4.**
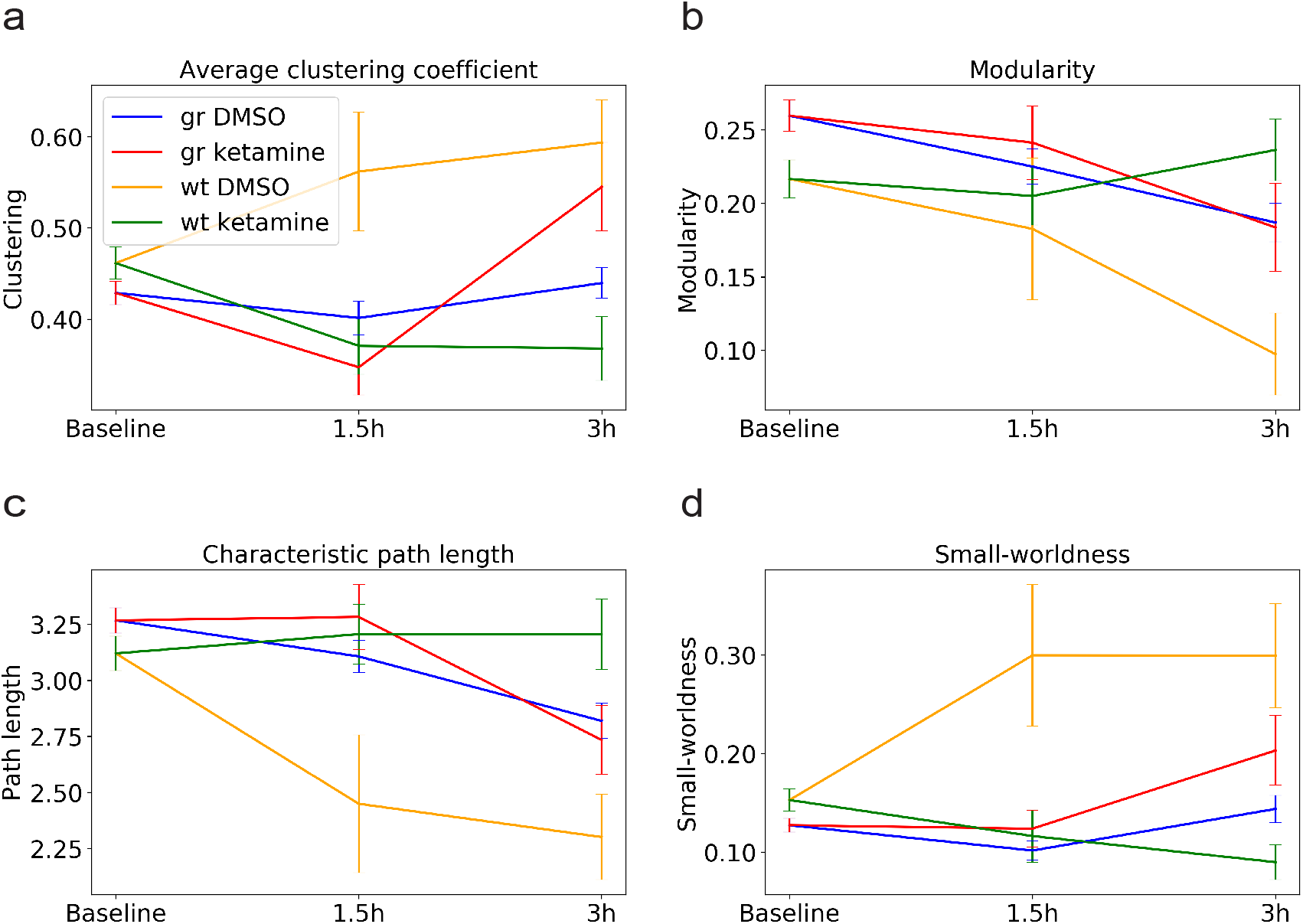
Effect of genotype and drug treatment with ketamine on network parameters. Mean of the **(a)** clustering coefficient, **(b)** modularity, **(c)** characteristic path length and **(d)** small-worldness in *gr^s357^* mutant and wildtype (wt) fish before drug treatment (baseline) and after 1.5h and 3h incubation with ketamine (20μM) or DMSO. 8 fish were imaged for each combination of genotype (*gr^s357^*, wt) and treatment (ketamine, control). Error bars represent the standard error of the mean.

Interestingly, ketamine differentially affected the two genotypes (Fig. 4). After incubation for 1.5h with ketamine, clustering (Fig. 4a; GR DMSO vs. GR ketamine: *p* = 0.019) was significantly reduced in *gr^s357^* mutants, while characteristic path length (Fig. 4c; GR DMSO vs. GR ketamine: *p* = 0.049) was increased. In wildtype, clustering (Fig. 4a; WT DMSO vs. WT ketamine: *p* = 0.006) and small-worldness (Fig. 4d; WT DMSO vs. WT ketamine: *p* = 0.003) were significantly reduced after incubation with ketamine compared to the wildtype DMSO control, while characteristic path length (Fig. 4c; WT DMSO vs. WT ketamine: *p* = 0.023) was increased. After 3h incubation with ketamine, we observed higher clustering (Fig. 4a; GR DMSO vs. GR ketamine: *p* = 0.026) and increased small-worldness (Fig. 4d; GR DMSO vs. GR ketamine: *p* = 0.043) in ketamine-treated *gr^s357^* mutant fish compared to the *gr^s357^* mutant DMSO control. In wildtype fish, differences appeared in all network parameters after 3h incubation with ketamine compared to wildtype DMSO control: Clustering (Fig. 4a; WT DMSO vs. WT ketamine: *p* = 0.001) and smallworldness (Fig. 4d; WT DMSO vs. WT ketamine: *p* = 0.000) were significantly reduced, whereas characteristic path length (Fig. 4c; WT DMSO vs. WT ketamine: *p* = 0.002) and modularity (Fig. 4b; WT DMSO vs. WT ketamine: *p* = 0.001) were increased. Thus, we observed a drug x genotype interaction effect (Table 1): Ketamine seems to increase functional connections and efficiency of global and local integration in the mutant. The opposite effect was observed in wildtype fish, where ketamine reduced both clustering and small-world properties of the neuronal network.

### Restoration of brain network function in *gr^s357^* mutants by cycloserine

Cycloserine is an NMDA receptor agonist at low and an NMDA antagonist at high dosages. After 1.5h incubation with cycloserine, statistical tests revealed that there was no significant difference in clustering (*p* = 0.160) (Fig. 5a), characteristic path length (*p* = 0.158) (Fig. 5c), modularity (*p* = 0.428) (Fig. 5b) or small-worldness (*p* = 0.087) (Fig. 5d) between cycloserine treated *gr^s357^* mutants and wildtype controls. After incubation for 3h, clustering was significantly lower in cycloserine treated *gr^s357^* fish compared to the wildtype DMSO control (*p* = 0.004); characteristic path length (*p* = 0.015) and modularity (*p* = 0.01) were increased, whereas small-worldness was reduced. These results suggest that cycloserine shows similar effects on network parameters as fluoxetine. Network parameters were not significantly different from wildtype control fish after application of the drug. However, this effect was not stable: after 3h of incubation, the normalizing effect of cycloserine disappears. Cycloserine dosage may be decreasing after 1.5h or may reach a concentration at which it acts as an NMDA receptor agonist; therefore, its antidepressant potential may be short-lived.

**Figure 5.**
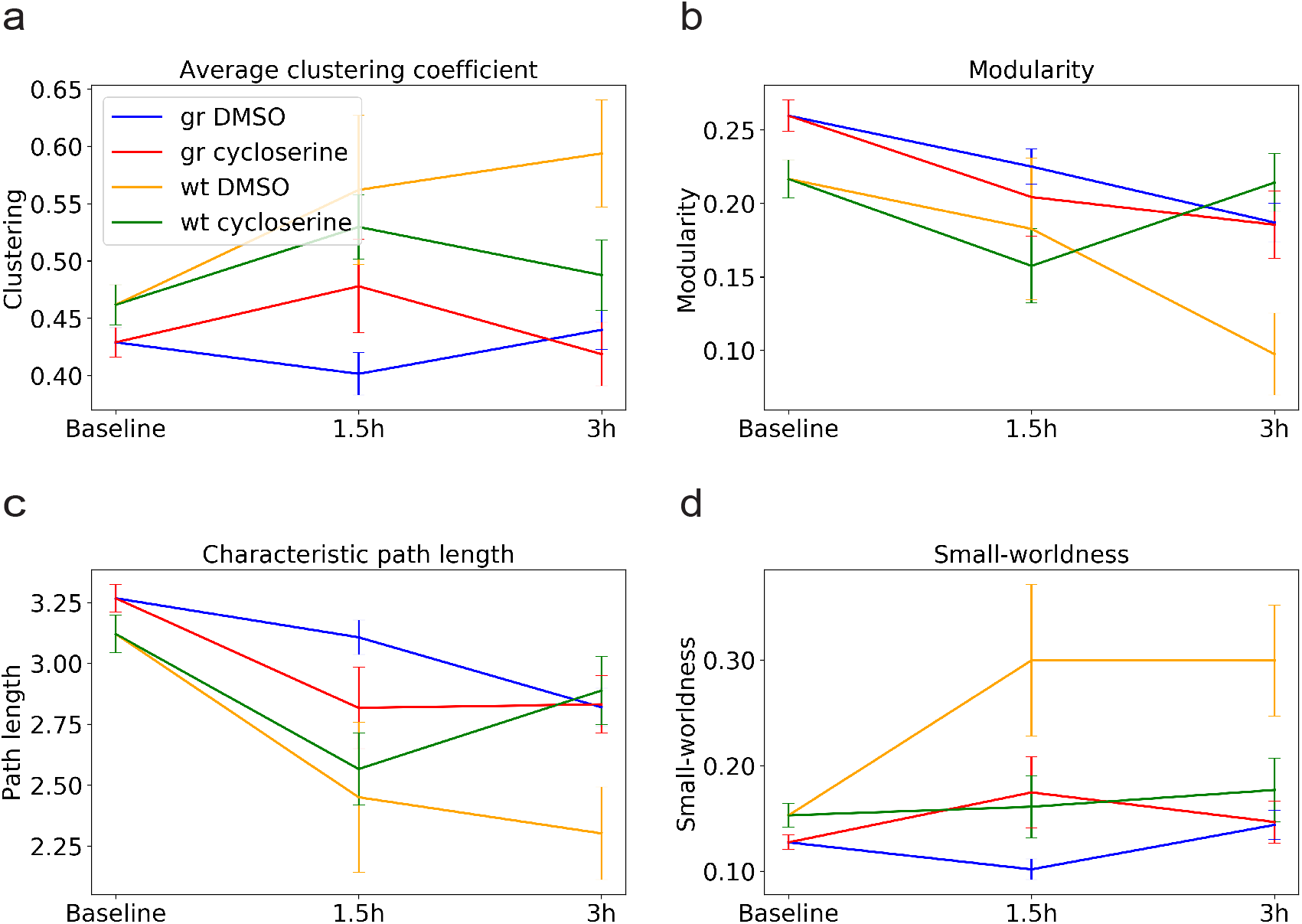
Effect of genotype and drug treatment with cycloserine on network parameters. Mean of the **(a)** clustering coefficient, **(b)** modularity, **(c)** characteristic path length and **(d)** small-worldness in *gr^s357^* mutant and wildtype (wt) fish before drug treatment (baseline) and after 1.5h and 3h incubation with cycloserine (20μM) or DMSO. 8 fish were imaged for each combination of genotype (*gr^s357^*, wt) and treatment (cycloserine, control). Error bars represent the standard error of the mean.

Cycloserine differentially affected *gr^s357^* mutants and wildtype. Incubation for 1.5h with cycloserine, increased clustering (Fig. 5a; GR DMSO vs. GR cycloserine: *p* = 0.028) and small-worldness (Fig. 5d; GR DMSO vs. GR cycloserine: *p* = 0.013) in mutants, but not in wildtype. This effect of cycloserine in mutants disappeared again after 3 hours, while clustering (Fig. 5a; WT DMSO vs. WT cycloserine: *p* = 0.020) and small-worldness (Fig. 5d; WT DMSO vs. WT cycloserine: *p* = 0.002) were reduced and modularity (Fig. 5b; WT DMSO vs. WT cycloserine: *p* = 0.003) as well as characteristic path length (Fig. 5c; WT DMSO vs. WT cycloserine: *p* = 0.013) were increased in wildtype. These results suggest that, for cycloserine, there is a drug x genotype interaction over the course of the 3h experiment (Table 1). We assume that, after incubation for 1.5h, cycloserine’s antagonistic effect is at its peak and decreasing inhibition in the mutant fish brain leads to a highly clustered functional network with augmented local and global efficiency in information processing and integration. In wildtype brains, we could not observe this effect. After 3h incubation, this effect disappeared in the mutant. For wildtype fish, functional connectivity is affected after 3h and network parameters are shifted, but interestingly in the opposite direction as in mutants. At this point, cycloserine may increase inhibition in the wildtype brain. Due to a potentially higher baseline level of inhibition, this effect is not present in the *gr^s357^* mutant fish.

### Conclusions and outlook

Taken together, our results demonstrate that graph analysis of light-sheet calcium imaging data is suited to reveal the global correlational structure of zebrafish whole-brain activity and has the sensitivity and resolution to detect changes caused by genetic or pharmacological perturbations. The construction of graphs from automatically segmented neurons allowed us to extract brain-wide network parameters. We propose that exploring drug effects and genotype x drug interactions in a computational pipeline that employs rapid light-sheet imaging followed by a first-pass graph analysis opens up new possibilities for understanding altered brain states in zebrafish larvae. Streamlining and further automating this pipeline may enable a small-molecule compound screen for the identification of novel antidepressants. In the future, this scalable approach can be applied to a variety of zebrafish disease models and tailored interventions in order to accelerate our understanding of disease conditions and their therapies.

## Methods

### Animal care and transgenic lines

All animal procedures conformed to the institutional guidelines of the Max Planck Society and the local government (Regierung von Oberbayern). Experimental protocols were approved by Regierung von Oberbayern (55.2-1-54-2532-41-2016 and 55.2-1-54-2532-31-2016). Fish were raised on a 14h light/10h dark cycle at 28°C according to standard procedures. Transgenic lines carrying the *gr^s357^* mutation were generated as described previously (Ziv et al., 2013), and for imaging purposes crossed to *Tg(elavl3:H2B-GCaMP6s)* obtained from Misha Ahrens (Janelia Research Campus).

### Light-sheet imaging

Larvae (*gr^s357/s357^* or *gr*^s357/+^; *Tg(elavl3:H2B-GCaMP6s)*) were pre-screened for expression of the genetically encoded calcium sensor GCaMP6s under the fluorescent microscope at 5 dpf. On the following day, fish were embedded in 1.5% low-melting-point agarose (Invitrogen) in a custom-designed, 3D-printed chamber and aligned to a template (see Fig. 1c, d, e). Once hardened, the block of agarose was cut to reduce scattering of the beam and immersed in Danieau’s solution to fill a total volume of 3 ml. Embedded larvae were allowed to accommodate in the setup for at least 30 minutes before the start of imaging.

The light-sheet microscope consists of two orthogonal excitation arms for frontal and lateral scanning, and a detection arm for imaging from above. A laser beam (473 nm wavelength, 3-5 mW power) is split by a dichroic mirror and guided to the excitation arms by six dielectric mirrors. Two pairs of mirror galvanometers oscillate horizontally and vertically to create a sheet of light (Fig. 1c) and scan the sample in z, respectively. Two serial lenses act as a pinhole and focus the sheet into the back aperture of a 4x objective. The detection arm consists of a 20x detection objective, a piezoelectric component to adjust the focus (10 V displacement, corresponds to a 400 μm range in z), a tube lens, fluorescence filter and an Orcaflash 4.0 sCMOS camera. The chamber containing the sample sits on a lab jack stage, which is brought into the field of view of the camera by x, y and z actuators.

Spontaneous brain activity of the larvae was imaged with a constant current of 35 mA, a frequency of 2 Hz and 60 frames per second at 6 ms exposure time, resulting in 30 planes in z, imaged twice per second (2 brain volumes per second) for a duration of 5 min in each condition. The ventral-dorsal range was specific to every larva, but usually ranged from 5 – 5.6 V, or 200 – 224 μm, respectively, resulting in a gap of approximately 7 μm between planes. To minimize file size, a summation of 4×4 pixels was applied, where one pixel corresponds to 330 nm (before summation) and 1.32 μm (after summation), respectively.

### Drug treatment

In three treatment conditions, the drug compounds (fluoxetine: Sigma-Aldrich, CAS No. 56296-78-7; ketamine: Sigma-Aldrich, CAS No. 1867-66-9; cycloserine: Sigma-Aldrich, CAS No. 68-41-7) were diluted in DMSO and applied in the setup to reach a final concentration of 20μM for each compound. In a fourth, control condition, fish larvae were incubated with DMSO only. Three light sheet recordings were acquired per fish, each lasting 5 min. The first recording was taken right before drug incubation started (baseline condition), and the other two were taken after 1.5 and 3 hours of incubation with the drug or control medium (see Fig. 1b). For image acquisition, genotypes and drug treatments were randomized. We recorded up to three fish in a day. Overall data from 64 fish were recorded, 8 for each combination of genotype (8 *gr^s357^* mutants, 8 wildtype) and 4 treatments (3 drugs, control).

### Analysis of functional imaging data

Four-dimensional imaging stacks (time, z, y, x) were split into time series of single planes and corrected for motion artifacts using an adapted version of the CaImAn package (Pnevmatikakis et al., 2017; Giovannucci et al., 2019). In brief, this consists of a rigid transformation, which grossly rotates the images in x and y, followed by an optic flow transformation for finer features. Individual neurons were detected using a procedure described previously (Portugues et al., 2014). In short, regions of interest (ROIs), predominantly corresponding to individual cells, were defined as clusters of co-active pixels, using time series correlations of each pixel with its neighboring pixels (see Fig. 1f, g). The algorithm picked up between 20,000 and 40,000 ROIs for each fish brain, of which 10,000 cells were selected randomly to construct the functional connectivity matrix used for graph analysis. Functional connectivity was computed as Pearson’s correlation coefficients between signal time series of these ROIs (see Fig. 1h). Using MATLAB (R2016b, The MathWorks, Natick, MA) and the Brain Connectivity Toolbox (Rubinov & Sporns, 2010), we then calculated the following graph-theoretic metrics: the clustering coefficient (reflecting functional segregation in the network), the characteristic path length (reflecting functional integration), modularity (reflecting the degree of segregation of the network into highly clustered modules), and small-worldness (reflecting an optimal balance of functional integration and segregation).

### Statistical analysis of network parameters

The Shapiro-Wilk test was used to test the null hypothesis that data for all variables originate from a normally distributed population. For further analysis non-parametric tests were used, because the test revealed that our data are not normally distributed. Network parameters including the clustering coefficient, modularity, characteristic path length and small-worldness were analyzed for differences between groups of genotypes and drug treatments at baseline and after 1.5h and 3h incubation with the Mann-Whitney U test using the Python module SciPy. To correct for multiple comparisons, we used Bonferroni correction. The *p* values quoted in the text are already adjusted, where appropriate, according to these corrections.

## Acknowledgments

We thank all members of our laboratory for discussions and support, especially Krasimir Slanchev. Misha Ahrens (Janelia Research Institute) provided the *elavl3:H2B-GCaMP6s* fish. The light-sheet microscope was built by Marco Dal Maschio and Joe Donovan, in collaboration with Ruben Portugues and his team (Max Planck Institute of Neurobiology) and with advice from Michael Orger (Champalimaud Institute, Lisbon).

## Author Contributions

J.B. and H.B. conceived and designed the project. J.B. and E.H. set up the light-sheet microscope and performed imaging experiments. J.B. performed graph analysis. B.G. provided MATLAB scripts. J.B., E.H. and H.B. wrote the manuscript.

## Notes

The authors declare that there are no potential conflicts of interest with respect to the research, authorship, and/or publication of this article.

#### Summary of Updates

Authors have been added.

